# Differential retention of Pfam domains creates long-term evolutionary trends

**DOI:** 10.1101/2022.10.27.514087

**Authors:** Jennifer James, Paul Nelson, Joanna Masel

## Abstract

Protein domains that emerged more recently in evolution have higher structural disorder and greater clustering of hydrophobic residues along the primary sequence. It is hard to explain how selection acting via descent with modification could act so slowly as not to saturate over the extraordinarily long timescales over which these trends persist. Here we hypothesize that the trends were created by a higher level of selection that differentially affects the retention probabilities of protein domains with different properties. This hypothesis predicts that loss rates should depend on disorder and clustering trait values. To test this, we inferred loss rates via maximum likelihood for animal Pfam domains, after first performing a set of stringent quality control methods to reduce annotation errors. Intermediate trait values, matching those of ancient domains, are associated with the lowest loss rates, making our results difficult to explain with reference to previously described homology detection biases. Simulations confirm that effect sizes are of the right magnitude to produce the observed long-term trends. Our results support the hypothesis that differential domain loss slowly weeds out those protein domains that have non-optimal levels of disorder and clustering. The same preferences also shape differential diversification of Pfam domains, further impacting proteome composition.

## Introduction

The possibility that evolution could result in long-term directional change over time has enduring appeal, but there are few well-documented examples. It has proven difficult to rigorously assess purported universal tendencies toward increases in complexity and size over time (Gregory 2008; McShea 1991). Perhaps the best documented example of a long-term trend is that some taxonomic groups show a tendency toward increasing body size over time, aka *‘*Cope*’*s rule*’* (Cope 1885; Heim et al. 2015; Payne et al. 2009). Interestingly, this trend does not seem to be caused by directional selection slowly exerting a preference for larger individuals, but rather by the differential diversification of larger body size clades (Heim et al. 2015). This is unsurprising, because directional selection among individuals of the same species works quickly, making it an unlikely explanation for such a slow directional trend.

New examples of long-term trends were recently observed in the properties of protein domains (e.g. intrinsic structural disorder) as a function of how much time has elapsed since their birth, with considerable work performed to exclude artifactual explanations (Foy et al. 2019; James et al. 2021). We find it more plausible that some bias in the evolutionary process is responsible for these trends, rather than that the external environments shaping birth have moved in such a consistent direction over such long timescales. One hypothesis is that regardless of time of birth, all domains were born with properties biased toward promoting *de novo* gene birth (Wilson et al. 2017), and since then have had different amounts of time to evolve away from that starting point, toward properties that are more optimal for correct folding (Bucciantini et al. 2002; Chiti and Dobson 2017; Debès et al. 2013). This hypothesis is supported by the fact that the amino acid frequencies characteristic of newborn animal domains also make the expression of a random peptide more benign in *Escherichia coli* (Kosinski et al. 2022). But it is very hard to explain why directional selection on amino acid frequencies has been so slow to take full effect. We know that the evolution of amino acid frequencies can be rapid, as it is able to keep pace with the rapid evolution of nucleotide composition (Brbić et al. 2015).

Here we consider the possibility that instead of being caused by bias during descent with modification, long-term trends as a function of age are due to the differential retention vs. loss of protein domains with different properties. We hypothesize that domains are born with a wide range of biochemical properties. From this initial diversity, “fitter” domains, i.e. those that duplicate more often and/or are lost less often, are more likely to have descendants in extant organisms, and thus to appear in the older domain age classes.

Under this hypothesis, we expect a protein property that shows a long-term trend with age to be predictive of loss rate. One observed trend, extending all the way back to the last universal common ancestor, is that younger domains tend to have *‘*clustered*’* rather than interspersed hydrophobic acids along their primary sequence (Foy et al. 2019; James et al. 2021). A second trend, restricted to the evolutionary history of animals, is that younger domains tend to have a higher level of intrinsic structural disorder (ISD), primarily due to trends in the frequencies of the 20 amino acids (James et al. 2021). Here we test whether ISD and hydrophobic clustering predict the loss rate of a domain.

What we are proposing is that the relevant level of selection may not be the one that selects among alternative alleles of the same gene (Fig. 1, top), but instead the level that selects among alternative domains capable of fulfilling the same cellular functions as genes duplicate, differentiate, and are lost (Fig.1, bottom). Rapid loss of recently born protein-coding genes has previously been postulated (Palmieri, Kosiol, and Schlötterer 2014; Schlötterer 2015; Tautz and Domazet-Lošo 2011). The number of domain copies also appears to be surprisingly changeable, with fairly extreme expansions and contractions observed even over short evolutionary timescales (Hahn, Han, and Han 2007; Moore and Bornberg-Bauer 2012), leaving plenty of scope for differential diversification and retention to be a biologically significant influence.

**Figure 1.**
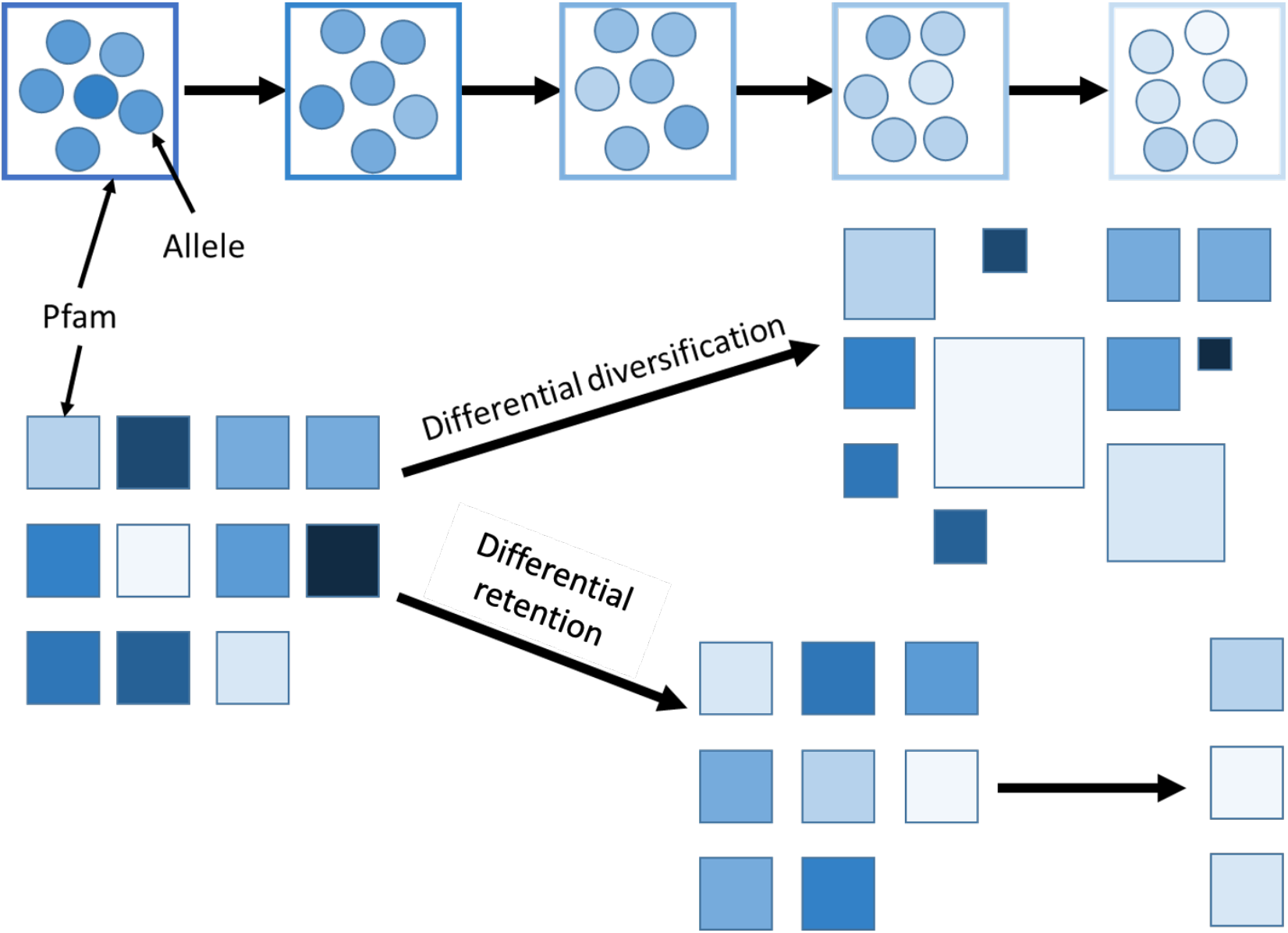
Alternative mechanisms/levels of directional selection over evolutionary time. Directional evolution in a trait with cohort age, represented by a change from dark to pale blue, can occur either via (top) descent with modification during which selected (pale blue) alleles replace other alleles, or (bottom) differential retention in which an initial diversity of Pfams is differentially retained. Differential diversification (middle) combines tendencies for duplication and tendencies for loss. Note that this is a levels of selection situation; a Pfam is a clade of protein-coding sequences and their component alleles, and the same figure could be interpreted with individual interests in lieu of allele interests, and clade interests in lieu of Pfam interests.

We focus on protein domains rather than on complete genes. We take domain annotations from the Pfam database (El-gebali et al. 2019), and refer to domains simply as Pfams. Protein-coding domains are sometimes thought of as functional units of protein sequence that are able to fold independently (Ponting and Russell 2002). However, Pfams are annotated by HMMER3 on the basis of evolutionary rather than functional relatedness (Finn, Clements, and Eddy 2011), and as such are fundamental units of protein sequence homology. This makes them easier to work with than genes, which commonly contain multiple different Pfams of a variety of ages (Bagowski, Bruins, and Velthuis 2010; Bornberg-Bauer and Albà 2013).

We infer loss rates by maximizing the likelihood of observed presence / absence of at least one Pfam instance across all species in a phylogeny. As well as this central focus on total loss rates, leading to differential retention over time (Fig. 1, bottom), we perform a complementary analysis of differential diversification (Fig. 1, middle) by taking the mean number of distinct instances of Pfams within a species. Both presence/absence data and number-of-instances data are vulnerable to both false positives and false negatives during homology detection. Our previous age assignment procedure (James et al. 2021) used a keyword based approach to remove Pfams that were likely incorrectly annotated in the focal genome due to contamination (Breitwieser et al. 2019; Lu and Salzberg 2018; Salzberg 2017), a problem that if uncorrected will result in nonsensical loss rates for affected Pfams. James et al. (2021) also removed Pfams that exhibited an implausible phylogenetic distribution, which could have resulted from either contamination or horizontal gene transfer. Here we go further, reducing the rate of false negatives by checking all six reading frames of contigs for unannotated Pfam instances, and then reconsidering whether the phylogenetic distribution of the Pfam is implausible. Second, we infer total loss rates jointly with a rate of false positive hits. Finally, we consider the hypothesis of homology detection bias while interpreting results.

We ask whether ISD and clustering scores predict the rate of total loss of a Pfam in a lineage, and also the number of Pfam instances per species. We analyse animal lineages only, because the ISD trends as a function of Pfam age found by (James et al. 2021) were specific to animals.

## Results

We used the phylostratigraphy dataset of James et al., restricted to the 6841 Pfams annotated in at least two out of our of 343 animal species, and already subject to a number of filters used by James et al. We scanned all six reading frames of intergenic regions to find species that contained unannotated instances of these Pfams. We also performed additional quality controls to remove likely viral or other contaminants on the basis of incoherent distribution across the tree (see Methods), resulting in a dataset of 6700 Pfams. We then estimated the rate of total loss of each Pfam over the animal species in our dataset (see Methods). During this maximum likelihood procedure, we iteratively removed Pfam presence data for clades (mostly single species) where they are more likely to be false positive hits or horizontal gene transfer events.

The median inferred loss rate of a Pfam in the animal phylogeny was 0.0009/MY (1^st^ and 3^rd^ quartiles of 0.00021 and 0.0035), i.e. 0.9 losses per 100,000 years. Note that this is not the rate at which a single instance of a Pfam is lost, but rather the rate at which all instances of a Pfam are lost from a lineage. Unsurprisingly, Pfams with a higher mean number of instances per genome tend to have a lower rate of total loss (Supplementary fig. 1, adjusted R^2^ = 0.30, p < 1e-16).

### Pfams with a mean ISD of 0.18 are least often lost

Pfams with high ISD have high loss rates (adjusted R^2^ = 0.023, slope = 1.35, *p* = 4e-36; data and loess regression shown in Fig. 2A). This is the predicted direction in order for their relationship to explain the animal phylostratigraphy trend (Fig. 3A). Note that the Box-Cox transformation of loss rates makes the units of the slope in these linear models hard to interpret. Interestingly, this relationship is not just non-linear, but also non-monotonic (loess regression in Fig. 2A). A regression model that includes a breakpoint at ISD = 0.18 (95% CI 0.15-0.21), representing the minimum loss rate, explains substantially more of the variance in loss rate among Pfams (adjusted R^2^ = 0.030, slopes -2.00 and 3.86), and fits the data significantly better than a model without a breakpoint (*p* = 6e-12), better than a model with zero slope for ISD values higher than the breakpoint (*p* = 1e-45), and better than a model with zero slope for ISD values lower than the breakpoint (*p* = 0.0002).

**Figure 2:**
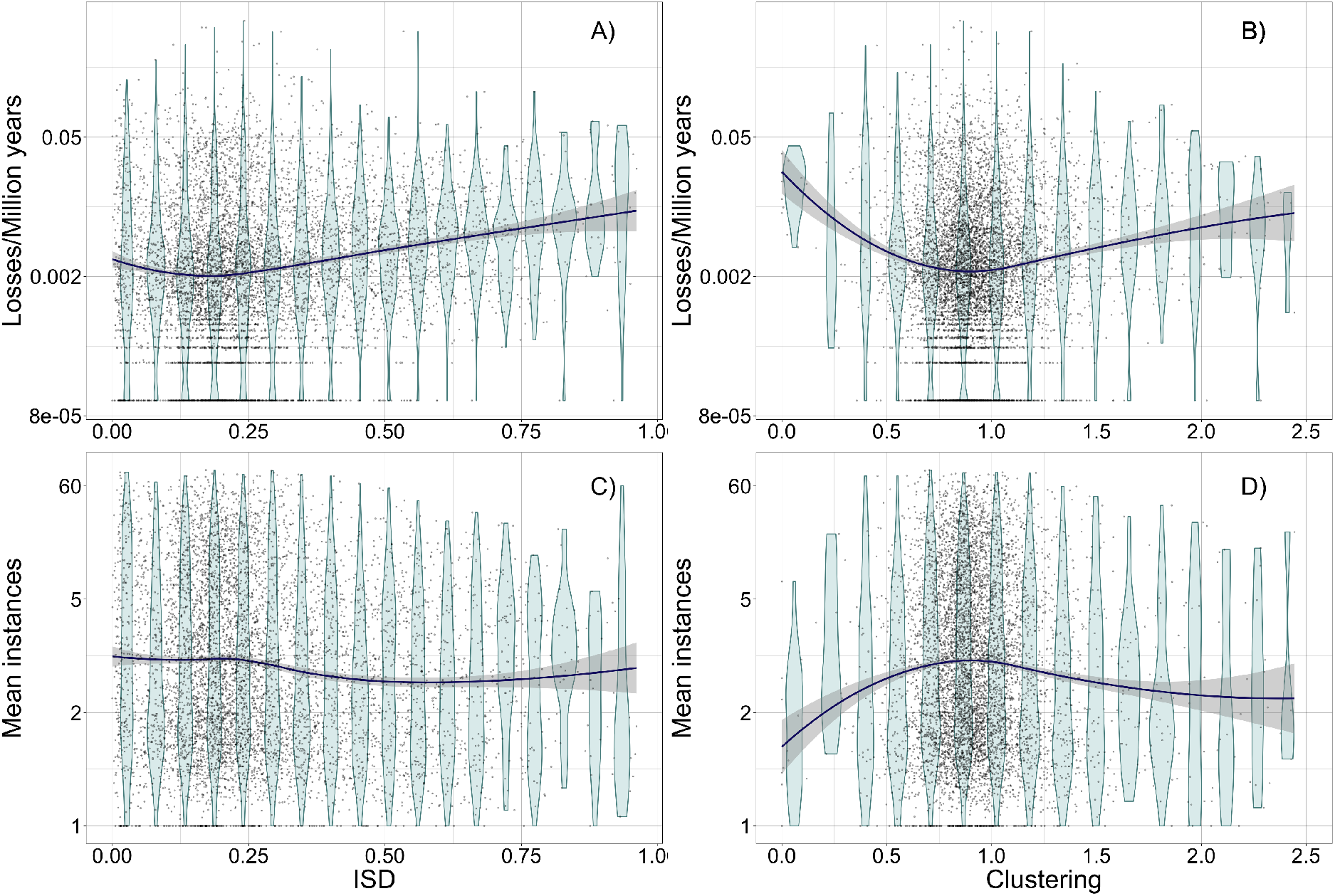
Pfam loss rates and copy number depend, often non-monotonically, on ISD and clustering values. The lowest loss rates are seen for Pfams with a mean ISD of 0.18 (A) and mean clustering value of 0.85 (B). Pfams with a low ISD have a higher number of instances per genome, in a more monotonic relationship (C). Pfams with a mean clustering value of 0.85 have the highest number of instances per genome. Each point represents a Pfam. Violin plots are a visual guide only, based on dividing the data into groups of equal range of ISD (A,C) or clustering (B, D). Loess regression curves are shown in dark blue, with 95% confidence intervals shown in grey. In A) and B), horizontal bands of points near the bottom represent 0, 1, 2, etc. observed loss events, while in C) and D) horizontal bands represent Pfams present in only a single copy per genome. In B) and D), for better visualization, the x-axis has been truncated at 2.5.

**Figure 3.**
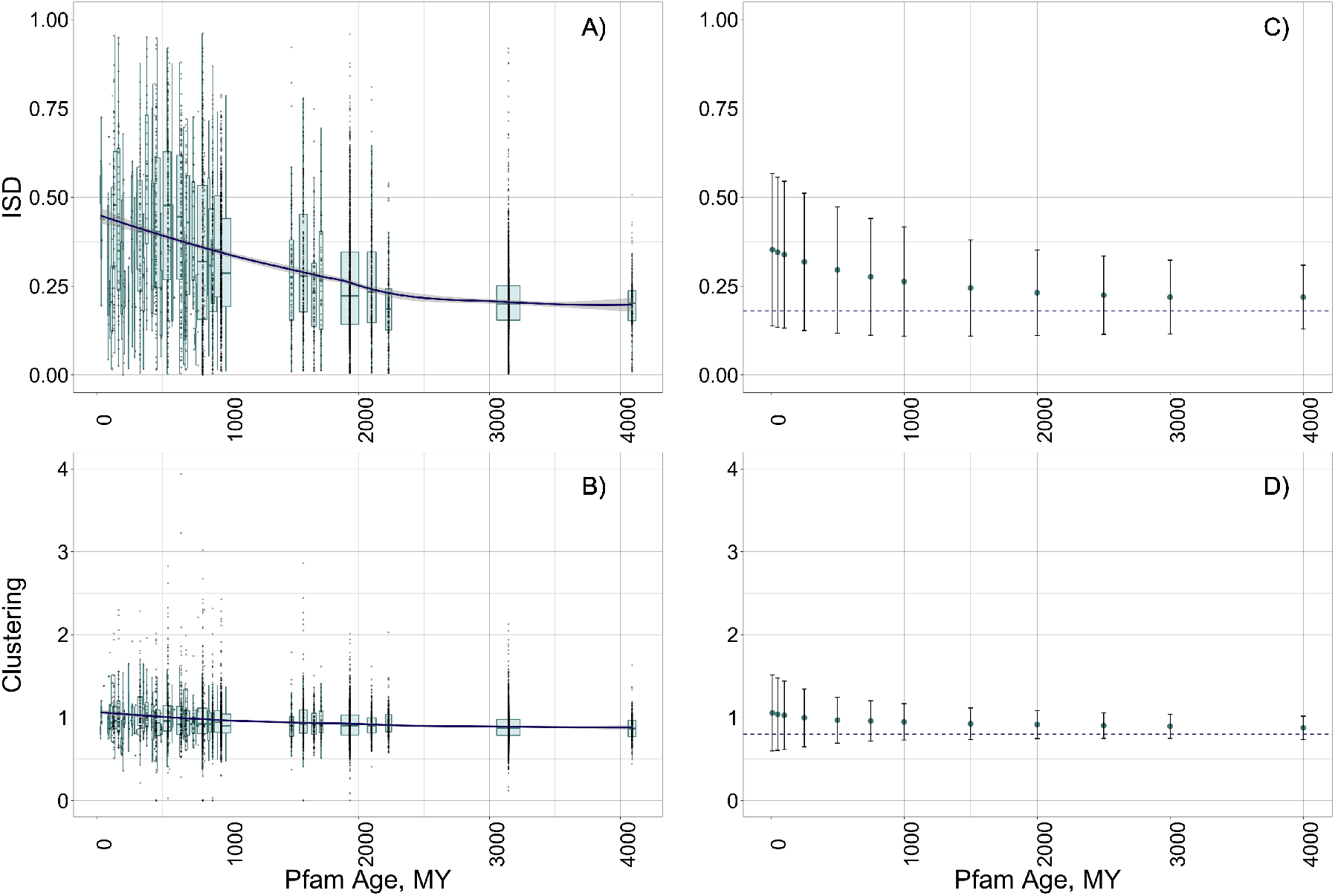
Observed phylostratigraphy trends in ISD (A) and Clustering (B) values occur on similar timescales as in our simulations of differential loss rates (C and D). In A) and B) each box represents all Pfams of a particular age category, with the widths of the boxes proportional to the square root of the number of Pfams in the group. Boxplots show the median, upper, and lower quartiles of the data, while the whiskers represent the largest and smallest data value above and below the interquartile range multiplied by 1.5. Grey points show the values for individual Pfams. The dark blue line is the loess regression slope of the effect of age on ISD (A) and clustering (B), with the 95% confidence interval shown in grey. In C and D, we plot the mean of ISD (C) and clustering (D), with error bars showing the standard deviation. Timepoints all use the same simulations and so are not independent, I.e. the number of domains in our simulations decreases with time. Dashed lines indicate the values of ISD and clustering corresponding to the lowest loss rates. Loss rates for the simulations are those inferred from piecewise linear regression from Figs. 2A and 2B.

The existence of a breakpoint rather than a monotonic relationship contradicts predictions from the homology detection bias hypothesis. That hypothesis expects overestimation of loss rates for high ISD Pfams whose homologs are more difficult to detect. While this could drive the overall relationship we found in which Pfams with high ISD have high loss rates, it cannot explain why loss rates also rise with low ISD <0.18.

Our results are instead consistent with selection acting to preferentially remove Pfams at either end of the spectrum of ISD. In further support of this, there is less variability in ISD within older Pfam cohorts (Fig. 4A, slope = -4.1-5 ISD/MY, adjusted R^2^ = 0.88, *p* = 3e-22, using linear regression weighted by number of Pfams per age cohort) than within younger Pfam cohorts. Ancient Pfams (those born over 2000MYA) that are still around today have a mean ISD of 0.21 (median 0.20), at the upper edge of the 95% confidence interval for the breakpoint representing minimum loss rate. These two observations further support the hypothesis that differential retention drives the long term trend in ISD as a function of evolutionary age observed in James et al (2021) and Foy et al. (2019). The fact that ancient Pfams may have ISD a little above the breakpoint might be noise in the data, or it might indicate that the trend of falling ISD with age (shown in Fig. 4A) has not yet fully saturated.

**Figure 4.**
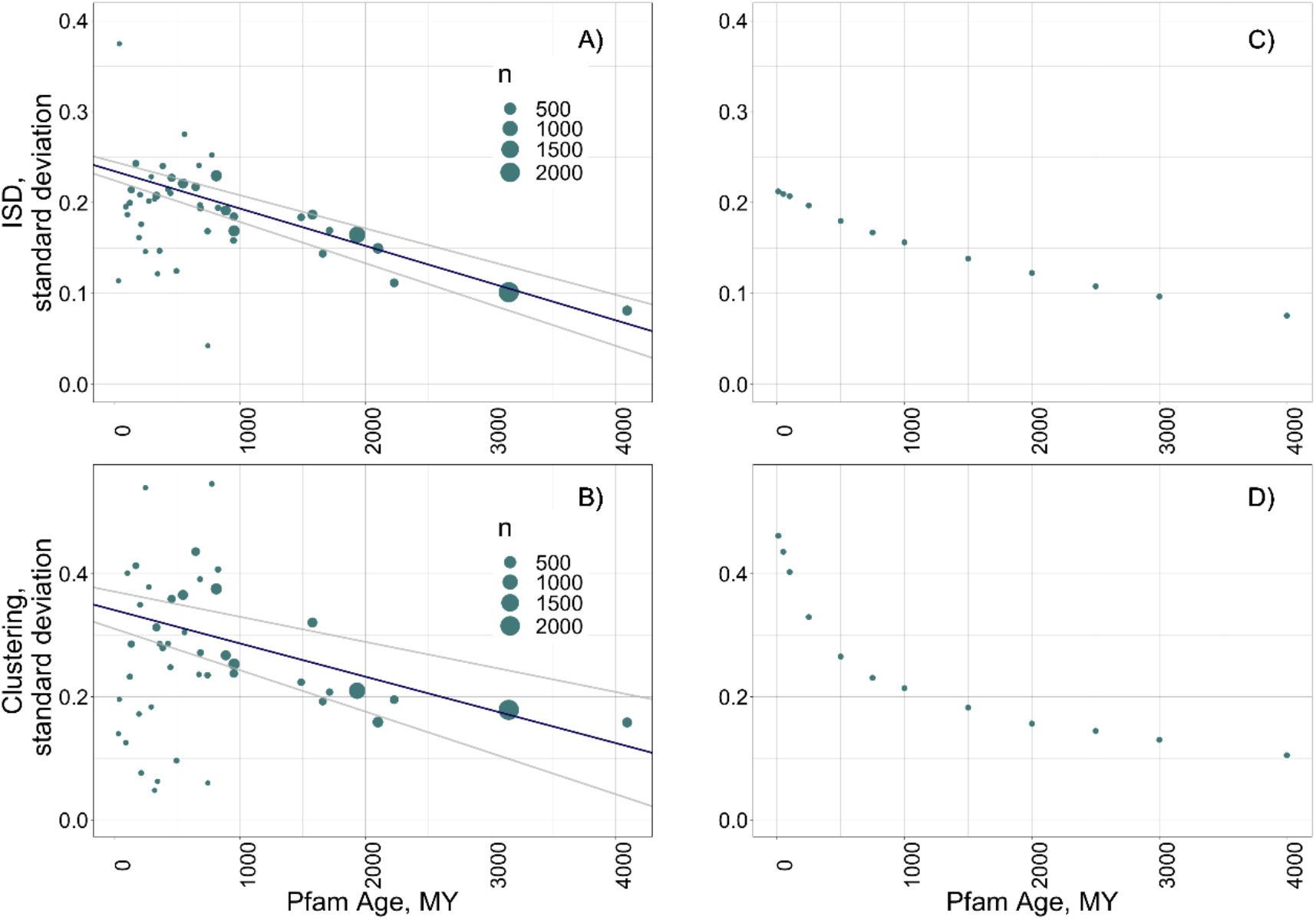
Younger Pfam cohorts have higher variance within their ISD (A) and clustering (B) distributions, a trend whose timescale is captured in our simulations (C, D). A) and B) Point size is scaled by n = number of Pfams in that category. Plotted in dark blue is the weighted linear regression slope (weighted by number of observations per category). Grey lines indicate the 95% confidence intervals on the regression. In A), for ISD, weighted R^2^ = 0.60, p = 2e-10. In B), for clustering, weighted R^2^ = 0.60, p = 2e-10. In C and D) the standard deviation of ISD (C) and clustering (D) values are plotted for cohorts of simulated Pfams.

### Pfams with moderately interspersed hydrophobic residues are least often lost

The hydrophobic residues of younger domains are more clustered (i.e. have higher clustering) than those of old domains (James et al. 2021). This directional trend could be explained mostly simply if Pfams with higher clustering were differentially lost. However, as with ISD, our data strongly support a non-monotonic relationship between clustering and loss rate (Figure 2B, losses/MY is two-parameter Box-Cox transformed prior to analyses). The best fitting linear model for clustering incorporated a breakpoint with a clustering value of 0.81 (95% confidence interval: 0.77 to 0.85) corresponding to the lowest loss rate. This model has an adjusted R^2^ of 0.020, and fits the data significantly better than a model without a breakpoint (*p* = 1e-25), than a model with zero slope for the lower clustering values (*p* = 8e-28), and than a model with zero slope for the higher clustering values (*p* = 2e-29). Clustering is not known to affect homology detection, but even if it did, it is again not clear how homology detection bias could produce a non-monotonic relationship.

A clustering value of 1 corresponds to a random distribution of hydrophobic residues, while the optimal clustering value of 0.81 represents interspersion of hydrophobic residues. Similar to the ISD results, this breakpoint value corresponds closely to the average clustering score of the most ancient Pfams (over 2000 MY old), which have mean and median clustering scores of 0.89 and 0.88, respectively. The fact that this is higher than the upper end of our 95% confidence interval for the breakpoint suggests that the trend in clustering values with age has not yet saturated. The y-intercept of Fig. 3B shows that Pfams are born with clustering scores a little above 1. Because Pfams tend to be born with relatively random amino acid placement corresponding to a clustering score of ∼1 that is so much higher than the optimum value of 0.81, differential Pfam retention can explain the progressive drop in clustering scores with Pfam age. In further support of the hypothesis that differential loss is responsible for the trend in clustering with age, there is more variability in clustering within younger Pfam cohorts (Fig. 4B, slope of standard deviation in clustering with age = -5.5 × 10^−5^, adjusted R^2^ = 0.60, p = 2e-10, using linear regression weighted by number of Pfams per age cohort).

### Results are robust to Pfam age as a confounding factor

When ascertaining the causal effects of ISD and clustering on loss rates, Pfam age has the potential to be a confounding factor in our analyses, for two reasons. Firstly, if older Pfams also have lower ISD and less clustered hydrophobic amino acids (James et al. 2021) for reasons other than differential loss due to these properties, this could potentially drive a spurious relationship between Pfam properties and loss rates. As expected, we observe lower loss rates for older Pfams in our data (Adjusted R^2^ = 0.050, *p* = 2e-76, supplementary figure 2). Secondly and more subtly, Pfams of identical age share the same speciation events, which enable Pfam losses to be observed. This results in phylogenetic structure in the data. A further complication is that the fact that older Pfams have survived to be observed might create ascertainment bias toward estimated loss rates that are lower than the true loss propensity.

To control for both indirect correlations and phylogenetic structure, we used mixed-effect linear regression models, including age as a random rather than a quantitative effect. Note that each branch of the tree corresponds to a possible Pfam age, making age a discrete quantitative variable. Empirically, we found that random effects models are better supported model choices: for example, for models of loss rate and ISD, R^2^ = 0.33 if we include age as a random effect, and R^2^ = 0.058 if we include age as a linear predictor, where R^2^ here is the conditional R^2^ describing the variance explained by the entire model. AIC scores for the respective models are 23580 and 24432. Similarly, for models of loss rate and clustering, a model that includes age as a random effect model has R^2^ = 0.34, while a fixed effect model has R^2^ = 0.05. AIC values are 23603 and 24476, respectively.

The non-monotonic relationship between loss rate and ISD remains statistically supported in the random effect model (*p* = 0.0003), with the model having a marginal R^2^, which represents the variance explained in loss rate just by the fixed effect of ISD, of 0.0030, and a slope of -1.17 for values of ISD below 0.18, and 0.64 for values above. The non-monotonic relationship with clustering also remained supported in the random effect model (*p* = 0.02), with a marginal R^2^ of 0.00054, with a slope of -0.24 for values of clustering below 0.81, and 0.097 for values of clustering above 0.81. While controlling for Pfam age causes marginal R^2^ values to drop dramatically, the causal direction is unclear. I.e., differences in ISD and clustering might drive changes in loss rates as a function of Pfam age, in which case controlling for age would remove a good portion of the genuine biological effect. However, the statistical significance we find indicates that Pfam age alone is insufficient to be the sole driver of the dependence of loss rates on ISD and clustering.

### Differences in loss rates are small enough to produce slow change

Even without controlling for phylogenetic structure/age, the effect sizes for loss and clustering on ISD are clearly small. In most contexts, this would cast doubt on their biological relevance. However, our motivating question is to discover what mechanism might have a small enough effect size such that its steady action could continue for billions of years before saturating. We therefore next use simulations to ask whether the effects of ISD and clustering on loss are of approximately the right magnitude to explain the relationship between these variables and Pfam age, as observed in James et al. (2021).

We simulate a set of Pfams initialized with the distribution of ISD or clustering values observed in young animal Pfams, and subject them to differential loss at rates given by our fitted regression models with breakpoints. This produces directional trends in ISD and clustering on the pertinent timescale (Fig. 3C, 3D). The relationship between ISD (Fig. 3A) and clustering (Fig. 3B) is visually extremely similar to that generated in our simulations (Figs. 3C and 3D). Our simple single-branch single loss event simulation ignores many complexities of the actual phylogenetic tree, but is sufficient to show that the effect sizes of ISD and clustering on loss rates are of approximately the right magnitude to explain phylostratigraphy trends.

### ISD and clustering also predict mean number of Pfam instances

The mean number of instances of the Pfam per genome is an alternative metric of evolutionary success. Not only is it likely an indirect proxy of total loss rate (supplementary fig. 1), but it also shapes proteome composition in its own right, independently of its effect on total loss rate (Figure 1, middle vs. bottom). The addition of substantially novel protein features through domain duplication followed by significant divergence (Nasir, Kim, and Caetano-Anollés 2014) is subsumed within our metric of instance number.

Unfortunately, mean number of instances per genome is expected to depend on genome annotation quality, which is likely to vary across the animal species in our dataset. We have at least partly addressed this problem by restricting analyses to genomes deemed “Complete”. In agreement with our loss rate results, Pfams with a higher mean number of instances per genome have lower levels of ISD (Figure 2C, R^2^ = 0.0069, *p* = 7e-11). Pfams with a higher mean number of instances per genome also have low levels of hydrophobic clustering (adjusted R^2^ = 0.0013, *p* = 0.0017). A non-linear model for the effect of ISD on mean number instances is only marginally supported over a linear model (*p* = 0.049 in model comparison with linear relationship, breakpoint = 0.57, 95% confidence intervals: 0.42, -.73), unlike the clear case for a non-linear relationship with loss rates. However, for hydrophobic clustering, we again find support for a non-linear relationship (Figure 2D, adjusted R^2^ = 0.0099), with a highly supported breakpoint at the same value of clustering (0.85, 95% confidence intervals: 0.79 - 0.90). A model with a clustering breakpoint fits the data significantly better than one without (*p* = 1e-13).

## Discussion

The persistence of trends in biochemical properties of protein domains (ISD and hydrophobic clustering) over such strikingly long timespans has been an unresolved puzzle. Our results suggest that at birth, Pfam domains have a wide range of properties, and are then differentially retained vs. lost as a function of these properties. Specifically, optimal retention is achieved with somewhat low ISD values close to 0.18, and with interspersion of hydrophobic amino acids corresponding to a clustering value of 0.8. The oldest cohort of Pfams has mean ISD and clustering close to these values, and there is less variance within old Pfam cohorts than within young Pfam cohorts. Simulations show that the effect sizes of differential retention as a function of ISD and clustering are of the right order of magnitude (i.e. small enough) to explain how phylostratigraphy trends can persist for such long evolutionary times without saturating. Conditional on retention, similar values of ISD and clustering also predict the mean number of domain instances per genome, magnifying the impact of these biases on proteome-wide composition.

This study implements high-throughput phylostratigraphy; while we implemented a number of novel measures to reduce the frequency of false positive and false negative homologs, our dataset will not be free of error. We focus on Pfam domains, for which homology detection is performed using the more sensitive program HMMER rather than BLASTP (El-gebali et al. 2019; Finn, Clements, and Eddy 2011). Concerns about homology detection bias usually focus on its potential for causing some rapidly evolving Pfams to be annotated as too young (Moyers and Zhang 2017; Weisman, Murray, and Eddy 2020). However, Pfam age is not a central focus of our analysis. The potential issue here is rather that homology detection bias might explain why Pfams with high ISD appear to be lost more often and have fewer instances per genome (a similar bias for clustering has not been documented). However, the known difficulty in detecting homology given higher ISD cannot explain our observed non-monotonic dependence of loss rates and instance number on ISD.

To detect as many true homologs as possible, instead of relying on existing genome annotations, we scan all six reading frames for hits. Each Pfam was as a result included in a mean of 7.35 and a median of 4 new animal species, out of 343. However, HMMER can produce false positives (Pearson, Li, and Lopez 2017); while use of curated Pfam seeds might reduce this problem, it will not completely eliminate it. We minimized impact by co-inferring false positive (or horizontally transferred) hits, with a higher rate for those found in our 6-frame scan, at the time of our inference of loss rate. Overall, these data quality measures resulted in a substantial decrease in the mean loss rate of approximately 14%. Another key quality control was to remove some Pfams altogether as potential contaminants not from the genomes under study. Our dataset and methodology represent a new standard for work on protein domain evolution, which, while imperfect, has resulted in a higher quality dataset for further analysis. Nevertheless, given the residual likely presence of false positive and false negative Pfam hits, we expect our results on total loss of a Pfam to be more robust to genome annotation issues than the number of Pfam instances per genome. However, we note that our instance number and loss rate results qualitatively agree.

While protein domains no doubt experience idiosyncratic selection pressures associated with their specific structures and functions within organisms, our findings suggest a more universal tendency in which Pfams with ISD and clustering values close to the *‘*optimum*’* are differentially retained in the long term. A candidate for a universal cause is that the same biophysical properties that promote correct folding also promote toxic aggregation, posing a universal dilemma for proteins (Bucciantini et al. 2002; Chiti and Dobson 2017). Foy et al. (2019) proposed that young genes tend to address this dilemma via a more “primitive” strategy, with high ISD representing a sacrifice of folding but safety from misfolding. The slow evolution of hydrophobic interspersion was proposed to eventually provide an alternative method for avoiding toxic aggregation that allows for tighter protein folds (Foy et al. 2019). Our findings suggest that the emergence of a tight protein fold might instead depend on good luck in the properties of the newborn protein sequence from which it derives. This idea is supported by lattice protein simulations designed to capture frustration between folding and aggregation, in which there is an abundance of lower fitness peaks, and where sequences born with high hydrophobicity (the lattice protein equivalent of low ISD) are more likely to find a high fitness peak (Bertram and Masel 2020).

The current study does not directly rule out directionality in descent with modification, which might be operating simultaneously, in either the same or the opposite direction to the forces documented here. While the evolution of amino acid frequencies can be rapid enough to keep pace with changes in nucleotide composition (Brbić et al. 2015), epistatic effects might lead to significant deceleration beyond a certain point (Vieira-Silva and Rocha 2008). Vertebrate species with higher effective population size tend to evolve higher ISD (Weibel et al. 2020), which is the opposite direction to what would be needed to produce the phylostratigraphy trends under the assumption that most ancestral lineages had high population size. While ISD was not directly assessed, long-term directional trends in amino acid frequencies via descent with modification have been inferred during early evolution (Groussin, Boussau, and Gouy 2013), as well as during the more recent evolution of cold tolerance (Fontanillas et al. 2017; Lecocq et al. 2021).

However, while selection among alternative alleles of the same domain (i.e. descent with modification) is often the default explanation for molecular evolution trends (fig. 1 top), we provide evidence for selection acting on a higher level, among protein domains, highlighting the importance of differential retention and diversification of domains as an evolutionary force in protein sequence evolution (fig. 1 middle and bottom). Selection acting on multiple different levels to affect a trait does not necessarily result in conflict between the levels (Lewontin 1970). This higher level is analogous to the dependence of speciation and extinction rates on the traits of species (see, for example (Aguilée et al. 2018)). An interesting parallel to our findings is the work of Heim et al. (2015), who found that long-term trends do not seem to be caused by directional selection, but rather by the differential diversification of certain clades. Here we find that the same broad category of explanation applies to long-term trends in protein properties as a function of age since they originated.

The study of long-term directional change in evolution, and the level of selection at which it occurs, has historically focused on species-level traits such as morphology. But there are several advantages of conducting such studies on proteins. Firstly, because Pfam domains are easily identified using HMMER, they are highly tractable for use in the study of higher levels of selection. In particular, it is easier to count how many instances there are of a Pfam in a genome than it is to identify all species in a clade.

Secondly, it is impossible to distinguish between e.g. high extinction rates vs. low speciation rates from observations of extant species distributions alone (Louca and Pennell 2020). An inherent difficulty is that we cannot observe extinct species once they are gone; fossil data is required to provide additional information (Mongiardino Koch, Garwood, and Parry 2021). However, when a domain undergoes complete loss in one lineage, we can directly infer that loss through comparison to other lineages in which it was not lost. Sophisticated methods developed to study speciation and extinction rates (Fitzjohn 2012) could be adapted to take advantage of this during the study of domain duplication and loss rates. Future studies could examine not just how loss rates depend on trait values, but also account for systematic variation in duplication and loss rates among clades, or over time. The study of protein domain diversification and loss is interesting not only to advance our understanding of the evolution of the proteome, but also as a powerful new study system for those interested more broadly in the causes of long-term directional trends in evolution, and in levels of selection.

## Methods

### Pfam selection

We begin with the Pfam phylostratigraphy dataset of James et al. (2021). Briefly, for a species to be included in this dataset, a “Complete” annotated genome had to be available from NCBI, and the species had to be present in Timetree (Hedges, Dudley, and Kumar 2006). We further restricted this dataset to those Pfams that were present in a minimum of 2 animal species, and for which both ISD and clustering scores could be calculated, resulting in a dataset of 6841 Pfams, and their distributions over a phylogenetic tree of 343 animal species. These Pfams had previously been subject to two quality control filtering steps: a keyword-based screening designed to exclude possible contaminants, and an additional z-score screen to exclude Pfams with species distributions indistinguishable from chance. Genes and Pfams annotated as mitochondrial were also already excluded from our starting database.

Pfam ages were initially taken from James et al. (2021), based on a species tree of 435 eukaryote species, with additional resolution among the two most ancient Pfam cohorts taken from (Weiss et al. 2016). While we calculate loss rates only within the animal species tree, our analysis includes Pfams that originated prior to animals and that are therefore also found in other taxa.

### False negatives

To detect potentially missing Pfam instances in unannotated genes, including those missed due to frameshift sequencing errors, we translated all regions not annotated as coding sequences (CDS) in all six frames. For this purpose, we downloaded annotated genomes of the 343 animal species in our dataset from NCBI, with access dates by species varying between May and July 2019. We removed sequences annotated as “CDS” and any sequences of “N” more than 3 nucleotides long. We then translated the resulting nucleic acid sequences in all six reading frames, not requiring a start codon for initiation but ending each polypeptide at stop codons, similar to the procedure implemented by Deutekom et al. (2019), and ran amino acid sequences through InterProScan version 5.33-72.0. We record all novel Pfam hits for each species. This resulted in each Pfam being included in a mean of 7.35 and a median of 4 new animal species. Although this six-frame search procedure generates false positives at an elevated rate relative to annotated Pfams, these false positive rates remain modest (see below). To avoid inflation of mean instances / genome by pseudogenes, we only correct putative false negatives in cases where the species in question would otherwise have zero instances of the Pfam.

### False positives

#### Excluding contaminant Pfams

As in James et al. (2021), we exclude any Pfam that was both present in fewer than half the species in the original dataset (i.e. not restricted to animals), and had a species distribution that was indistinguishable from chance. Such Pfams are likely to be contaminants. Briefly, for each number of species *n*, we performed 20,000 simulations in which we placed Pfams in *n* randomly selected species. We then inferred the number of total losses by parsimony, and used the mean and variance for the resulting distribution of simulated numbers of total losses to calculate *z* scores for each empirically observed distribution of *n* Pfam instances. To remove Pfams with species distributions indistinguishable from chance, James et al. excluded Pfams with *z* scores < -2. Here, we used an additional screen where we excluded any Pfams that had *z* scores < -2 either with or without the intergenic hits flagged as possible false negatives as described above. This resulted in the further removal of 141 Pfams, leaving a dataset of 6700 Pfams.

#### Excluding false positive Pfam presence while estimating loss rates by maximum likelihood

We jointly estimate the total loss rate of each Pfam by maximum likelihood, together with the identity of false positive or horizontally transferred hits/instances. First consider the case of no false positive or horizontally transferred hits. Using branch lengths obtained from Timetree, we assume that loss events are independent, making the probability of loss of Pfam *i* with loss rate *λi* per million years on branch *j* of length *t*_*j*_ equal to*1* − *exp*(−*t*_*i*_*λ*_*j*_). The likelihood that Pfam *i* is completely lost on branches *j* and retained in at least one copy on branches *k*, given total loss rate *λ*_*i*_, is:

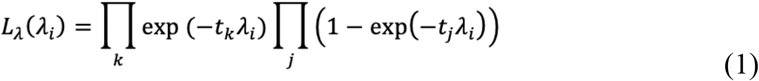

We first provisionally assign losses by assuming a Pfam arose on the branch basal to the monophyletic clade that includes all species with the Pfam, and is lost on the branches that explain the current distribution with fewest number of losses, i.e. we use Dollo parsimony (see figure 5). We then use Newton*’*s method to find the loss rate that maximizes the log likelihood, initialized at the loss rate calculated as the number of losses/total branch length of the Pfam tree. Because multiple loss events can occur on the same branch, the estimated ML loss rate estimate is generally very close but not identical to the initialization value (parsimony mean and median: 0.0058, 0.00088, ML mean and median: 0.0067, 0.00094). Occasionally, false positives / horizontal transfer events that are not yet accounted for trigger runaway inference of absurdly high ML loss rates at this stage. When the ML loss rate > 1 per million years, we therefore reset the loss rate to the initialization value originally estimated under parsimony, with associated likelihood, prior to proceeding.

**Figure 5.**
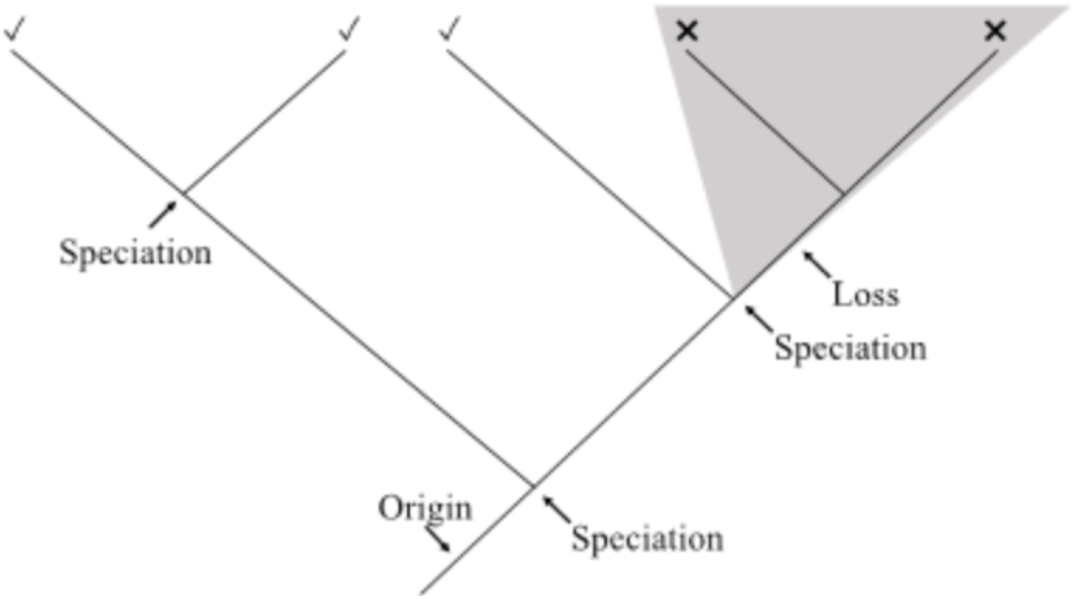
Inferring loss events for a Pfam by Dollo parsimony. Dollo parsimony assumes a Pfam arose exactly once, on the branch basal to the monophyletic clade that includes all species with the Pfam, and is lost on the branches that explain the current distribution with the fewest number of losses. Species with at least one instance of the Pfam are indicated with checks, and those without by crosses. We initialize the Pfam loss rate as 1/sum of unshaded branch lengths, i.e. all unshaded branches are counted by index *k* while the most basal shaded branch is counted by index *j* in Equation (1). We then refine this coarse Dollo-parsimony based estimate through maximum likelihood techniques that allow for false positive/horizontally transferred instances (Equation 2).

Next, to remove false positive / horizontally transferred Pfam presence data, we use a recursive empirical Bayesian procedure. We calculated two prior probabilities: *f*_*c*_, that the annotated presence of a Pfam in a species or clade is a false positive, and *f*_*i*_, that a Pfam presence discovered only by our six reading frame scan of intergenic sequences is a false positive. These were initialized at 10^−6^ and 0.2, respectively.

If *P*_*C*_ is the set of species annotated as having the Pfam from CDS observations, and *P*_*i*_ is the set of species annotated as having the Pfam only by us from the InterProScan intergenic search, the likelihood of observing the data if the species sets *F*_*C*_ and *F*_*i*_ are false positives is:

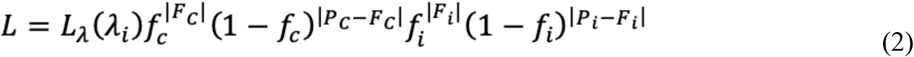

where *λ*_*i*_ is re-estimated by maximum likelihood for each species set *F*_*C*_ and *F*_*i*_, and absolute magnitude notation indicates the number of species in the set.

To find the sets *F*_*C*_ and *F*_*i*_ that maximize likelihood, we first group Pfam presence observations into the largest possible monophyletic clades where all species putatively possess at least one copy of the Pfam of interest. We then begin by “flipping” each such monophyletic clade, i.e. designating all species in a clade as being false positive hits or horizontal transfer events, and determining whether the likelihood improves. The false positive designation that improves the likelihood the most is kept, and then the process is repeated. In these subsequent iterations, we also test whether flipping a false positive back to a true positive improves likelihood. The process ends when no single flip improves the likelihood. After applying this process to all Pfams, we use the resulting set of false positive exclusion to estimate the two false positive probabilities. Using this as a revised empirical prior, we then repeat the procedure until no further change in the false positive rates is observed. Our method rapidly converges within 4 iterations, to a false positive rate of 0.00021 for Pfams previously annotated as coding, and 0.031 for Pfams annotated here by us based on presence in putatively intergenic regions. We calculate the mean number of Pfam instances per animal species within the set of species annotated as true positives at the end of the procedure described above.

Our procedures to include presence data when Pfams were previously unannotated, and to exclude presence data as described above, had a noticeable quantitative impact on our data. The combined effect of these quality control measures caused the mean and median loss rate to fall from 0.0075 and 0.00091 to 0.0065 and 0.00090. The mean and median number of Pfam instances per species rose from 5.9 and 2.2 to 6.2 and 2.4.

For some young Pfams (appearing after divergence with plants), our procedures to reduce false negative and false positive errors in presence annotation within the animal species tree lead us to re-estimate Pfam age. All of the Pfam loss rate and age data is available online at: https://github.com/j-e-james/DomainLossRate.

### ISD

We calculated predicted ISD using IUPred 2 (Dosztányi et al. 2005; Mészáros, Erdös, and Dosztányi 2018), which scores each amino acid between 0 and 1, with a low score indicating low disorder. In order to obtain a single estimate for each specific Pfam instance, we averaged over all amino acids. Per Pfam scores were then calculating by finding the mean over all copies of the Pfam in our dataset (i.e. animal species with more Pfam instances had higher weight).

### Clustering

Briefly, we calculated clustering as a kind of normalized index of dispersion: the variance in the number of the most hydrophobic amino acids (Leu, Ile, Val, Phe, Met and Trp) among blocks of 6 consecutive amino acids compared to the mean number, normalized so as to be comparable across Pfams of different lengths and hydrophobicities. If the length of a Pfam was not a multiple of 6, we took the average of all possible amino acid frames of 6, truncating the ends appropriately. For more detail, see Foy et al (2019) and James et al (2021). A clustering value of 1 indicates random distribution of hydrophobic residues, with higher values indicating that hydrophobic residues tend to be arranged in clusters along the sequence, and lower values indicating that hydrophobic residues are more evenly distributed along the sequence than would be expected by chance. As for ISD, per Pfam scores were then calculated as the mean over all instances of each Pfam in our animal dataset.

### Statistics

All plotting and statistical analysis was performed in R. Linear regression models that incorporate breakpoints were implemented using the segmented package. We used the packages lme4, MuMIn, and segmented to conduct models, and ggplot2 to create plots. Prior to linear regression, we transformed loss rate using an optimized two parameter Box-Cox transform (*λ* = 0.0054, *λ*^2^ = 6.1e-06) calculated using the R package geoR. We transformed mean instance number using a Box-Cox transform (*λ*= -0.61). We transform after rather than before taking the mean because the natural units for scoring impact on the proteome (Fig. 1 middle) are linear. Histograms comparing the distribution of untransformed versus transformed loss rates, in addition to Q-Q plots illustrating the requirement for transformation in our linear regression models with ISD, are shown in supplementary figure 3.

Throughout this work, we report the adjusted R^2^ values for our linear regression model results, as implemented in the programming language R, which uses the Wherry formula (Wherry 1931). This R^2^ accounts for the number of explanatory variables included in the model. Nested models were compared using the anova function in base R. We conducted mixed effects models in R using the lmer package, and calculated corresponding pseudo R^2^ values using the R package MumIn. This package returns marginal R^2^ values, which can be interpreted as the variance explained by fixed effects in the model, and conditional R^2^ values, which can be interpreted as the variance explained by both the fixed and random effects in the model (Nakagawa and Schielzeth 2013). R scripts and data required to replicate our analysis are available at: https://github.com/j-e-james/DomainLossRate.

### Simulations

We performed simulations to assess the biological relevance of our inferred loss rates, using custom python code available at https://github.com/j-e-james/DomainLossRate. We initialize these simulations by sampling, with replacement, the Pfams that we estimate to be younger than 100 MY old, to produce an initial set of 5000 Pfams, each represented by its ISD and clustering values. We then assign each Pfam the loss rate predicted from either its ISD value, or its clustering value, on the basis of our corresponding regression model. For each Pfam, we then sample the time to a single loss from an exponential distribution, with scale parameter = 1/loss rate. We then track the mean and standard deviation of the population of survivors over time.

## Acknowledgements

We thank Cecile Ané for helpful discussions that prompted a more complex project that we abandoned in favor of this simpler approach. We thank Nathan Aviles for helpful discussions about fancier statistical approaches that we didn*’*t use. We thank all of the participants at the 2019 “Workshop on Long-Term Trends in Evolution” for stimulating discussions. Both this work and the workshop were primarily supported by the John Templeton Foundation (60814). This work was also supported by the National Institutes of Health (GM-104040), and in its final stages, by the John Templeton Foundation (62220) and a Wenner-Gren postdoctoral stipendium.

## References

Aguilée, Robin, Fanny Gascuel, Amaury Lambert, and Regis Ferriere. 2018. “Clade Diversification Dynamics and the Biotic and Abiotic Controls of Speciation and Extinction Rates.” Nature Communications 9 (1): 1–13. https://doi.org/10.1038/s41467-018-05419-7.

Bagowski, Christoph P, Wouter Bruins, and Aartjan J W Velthuis. 2010. “The Nature of Protein Domain Evolution : Shaping the Interaction Network.” Current Genomics 11: 368–76.

Bertram, Jason, and Joanna Masel. 2020. “Evolution Rapidly Optimizes Stability and Aggregation in Lattice Proteins despite Pervasive Landscape Valleys and Mazes.” Genetics. https://doi.org/10.1101/776450.

Bornberg-Bauer, Erich, and M. Mar Albà. 2013. “Dynamics and Adaptive Benefits of Modular Protein Evolution.” Current Opinion in Structural Biology 23 (3): 459–66. https://doi.org/10.1016/j.sbi.2013.02.012.

Brbić, Maria, Tobias Warnecke, Anita Kriško, and Fran Supek. 2015. “Global Shifts in Genome and Proteome Composition Are Very Tightly Coupled.” Genome Biology and Evolution 7 (6): 1519– 32. https://doi.org/10.1093/gbe/evv088.

Breitwieser, Florian P., Mihaela Pertea, Aleksey V. Zimin, and Steven L. Salzberg. 2019. “Human Contamination in Bacterial Genomes Has Created Thousands of Spurious Proteins.” Genome Research 29 (6): 954–60. https://doi.org/10.1101/gr.245373.118.

Bucciantini, Monica, Elisa Giannoni, Fabrizio Chiti, Fabiana Baroni, Niccolò Taddei, Giampietro Ramponi, Christopher M. Dobson, and Massimo Stefani. 2002. “Inherent Toxicity of Aggregates Implies a Common Mechanism for Protein Misfolding Diseases.” Nature 416 (6880): 507–11. https://doi.org/10.1038/416507a.

Chiti, Fabrizio, and Christopher M. Dobson. 2017. “Protein Misfolding, Amyloid Formation, and Human Disease: A Summary of Progress Over the Last Decade.” Annual Review of Biochemistry 86 (1): 27–68. https://doi.org/10.1146/annurev-biochem-061516-045115.

Cope, E. D. 1885. “On the Evolution of the Vertebrata, Progressive and Retrogressive.” The American Naturalist 19 (2): 140–48.

Debès, Cédric, Minglei Wang, Gustavo Caetano-Anollés, and Frauke Gräter. 2013. “Evolutionary Optimization of Protein Folding.” PLoS Computational Biology 9 (1): e1002861. https://doi.org/10.1371/Citation.

Deutekom, Eva S., Julian Vosseberg, Teunis J.P. Van Dam, and Berend Snel. 2019. “Measuring the Impact of Gene Prediction on Gene Loss Estimates in Eukaryotes by Quantifying Falsely Inferred Absences.” PLoS Computational Biology 15 (8): 1–15. https://doi.org/10.1371/journal.pcbi.1007301.

Dosztányi, Zsuzsanna, Veronika Csizmók, Péter Tompa, and István Simon. 2005. “The Pairwise Energy Content Estimated from Amino Acid Composition Discriminates between Folded and Intrinsically Unstructured Proteins.” Journal of Molecular Biology 347 (4): 827–39. https://doi.org/10.1016/j.jmb.2005.01.071.

El-gebali, Sara, Jaina Mistry, Alex Bateman, Sean R Eddy, Simon C Potter, Matloob Qureshi, Lorna J Richardson, et al. 2019. “The Pfam Protein Families Database in 2019” 47 (October 2018): 427– 32. https://doi.org/10.1093/nar/gky995.

Finn, Robert D., Jody Clements, and Sean R. Eddy. 2011. “HMMER Web Server: Interactive Sequence Similarity Searching.” Nucleic Acids Research 39 (Web Server issue): 29–37. https://doi.org/10.1093/nar/gkr367.

Fitzjohn, Richard G. 2012. “Diversitree: Comparative Phylogenetic Analyses of Diversification in R.” Methods in Ecology and Evolution 3 (6): 1084–92. https://doi.org/10.1111/j.2041-210X.2012.00234.x.

Fontanillas, Eric, Oxana V. Galzitskaya, Odile Lecompte, Mikhail Y. Lobanov, Arnaud Tanguy, Jean Mary, Peter R. Girguis, Stéphane Hourdez, and Didier Jollivet. 2017. “Proteome Evolution of Deep-Sea Hydrothermal Vent Alvinellid Polychaetes Supports the Ancestry of Thermophily and Subsequent Adaptation to Cold in Some Lineages.” Genome Biology and Evolution 9 (2): 279– 96. https://doi.org/10.1093/gbe/evw298.

Foy, Scott G, Benjamin A Wilson, Jason Bertram, Matthew H J Cordes, and Joanna Masel. 2019. “A Shift in Aggregation Avoidance Strategy Marks a Long-Term Direction to Protein Evolution.” Genetics 211 (April): 1345–55. https://doi.org/10.1534/genetics.118.301719.

Gregory, T. Ryan. 2008. “Evolutionary Trends.” Evolution: Education and Outreach 1 (3): 259–73. https://doi.org/10.1007/s12052-008-0055-6.

Groussin, M., B. Boussau, and M. Gouy. 2013. “A Branch-Heterogeneous Model of Protein Evolution for Efficient Inference of Ancestral Sequences.” Systematic Biology 62 (4): 523–38. https://doi.org/10.1093/sysbio/syt016.

Hahn, Matthew W., Mira V. Han, and Sang Gook Han. 2007. “Gene Family Evolution across 12 Drosophila Genomes.” PLoS Genetics 3 (11): 2135–46. https://doi.org/10.1371/journal.pgen.0030197.

Hedges, S. Blair, Joel Dudley, and Sudhir Kumar. 2006. “TimeTree: A Public Knowledge-Base of Divergence Times among Organisms.” Bioinformatics 22 (23): 2971–72. https://doi.org/10.1093/bioinformatics/btl505.

Heim, Noel A., Matthew L. Knope, Ellen K. Schaal, Steve C. Wang, and Jonathan L. Payne. 2015. “Cope’s Rule in the Evolution of Marine Animals.” Science 347 (6224): 867–70. https://doi.org/10.1126/science.1260065.

James, Jennifer E., Sara M. Willis, Paul G. Nelson, Catherine Weibel, Luke J. Kosinski, and Joanna Masel. 2021. “Universal and Taxon-Specific Trends in Protein Sequences as a Function of Age.” ELife 10: 1–23. https://doi.org/10.7554/eLife.57347.

Kosinski, Luke J, Nathan R Aviles, Kevin Gomez, and Joanna Masel. 2022. “Random Peptides Rich in Small and Disorder-Promoting Amino Acids Are Less Likely to Be Harmful.” Genome Biology and Evolution 14 (6): 1–14. https://doi.org/10.1093/gbe/evac085.

Lecocq, Michel, Mathieu Groussin, Manolo Gouy, and Céline Brochier-Armanet. 2021. “The Molecular Determinants of Thermoadaptation: Methanococcales as a Case Study.” Molecular Biology and Evolution 38 (5): 1761–76. https://doi.org/10.1093/molbev/msaa312.

Lewontin, R. C. 1970. “Units of Selection.” Annual Review of Ecology and Systematics 1: 1–18. https://doi.org/10.1016/B978-008045405-4.00810-7.

Louca, Stilianos, and Matthew W. Pennell. 2020. “Extant Timetrees Are Consistent with a Myriad of Diversification Histories.” Nature 580 (7804): 502–5. https://doi.org/10.1038/s41586-020-2176-1.

Lu, Jennifer, and Steven L. Salzberg. 2018. “Removing Contaminants from Databases of Draft Genomes.” PLoS Computational Biology 14 (6): 1–13. https://doi.org/10.1371/journal.pcbi.1006277.

McShea, Daniel W. 1991. “Complexity and Evolution: What Everybody Knows.” Biology and Philosophy 6 (3): 303–24. https://doi.org/10.1007/BF00132234.

Mészáros, Bálint, Gábor Erdös, and Zsuzsanna Dosztányi. 2018. “IUPred2A: Context-Dependent Prediction of Protein Disorder as a Function of Redox State and Protein Binding.” Nucleic Acids Research 46 (Web Server issue): W329–37. https://doi.org/10.1093/nar/gky384.

Mongiardino Koch, Nicolás, Russell J. Garwood, and Luke A. Parry. 2021. “Fossils Improve Phylogenetic Analyses of Morphological Characters.” Proceedings of the Royal Society B: Biological Sciences 288 (1950). https://doi.org/10.1098/rspb.2021.0044.

Moore, Andrew D., and Erich Bornberg-Bauer. 2012. “The Dynamics and Evolutionary Potential of Domain Loss and Emergence.” Molecular Biology and Evolution 29 (2): 787–96. https://doi.org/10.1093/molbev/msr250.

Moyers, Bryan A., and Jianzhi Zhang. 2017. “Further Simulations and Analyses Demonstrate Open Problems of Phylostratigraphy.” Genome Biology and Evolution 9 (6): 1519–27. https://doi.org/10.1093/gbe/evx109.

Nakagawa, Shinichi, and Holger Schielzeth. 2013. “A General and Simple Method for Obtaining R2 from Generalized Linear Mixed-Effects Models.” Methods in Ecology and Evolution 4 (2): 133– 42. https://doi.org/10.1111/j.2041-210x.2012.00261.x.

Nasir, Arshan, Kyung Mo Kim, and Gustavo Caetano-Anollés. 2014. “Global Patterns of Protein Domain Gain and Loss in Superkingdoms.” PLoS Computational Biology 10 (1). https://doi.org/10.1371/journal.pcbi.1003452.

Palmieri, Nicola, Carolin Kosiol, and Christian Schlötterer. 2014. “The Life Cycle of Drosophila Orphan Genes.” ELife 3: 1–21. https://doi.org/10.7554/elife.01311.

Payne, Jonathan L., Alison G. Boyer, James H. Brown, Seth Finnegan, Micha Kowalewski, Richard A. Krause, S. Kathleen Lyons, et al. 2009. “Two-Phase Increase in the Maximum Size of Life over 3.5 Billion Years Reflects Biological Innovation and Environmental Opportunity.” Proceedings of the National Academy of Sciences of the United States of America 106 (1): 24– 27. https://doi.org/10.1073/pnas.0806314106.

Pearson, William R., Weizhong Li, and Rodrigo Lopez. 2017. “Query-Seeded Iterative Sequence Similarity Searching Improves Selectivity 5 – 20-Fold.” Nucleic Acids Research 45 (7): e46. https://doi.org/10.1093/nar/gkw1207.

Ponting, Chris P., and Robert R. Russell. 2002. “The Natural History of Protein Domains.” Annual Review of Biophysics and Biomolecular Structure 31 (1): 45–71. https://doi.org/10.1146/annurev.biophys.31.082901.134314.

Salzberg, Steven L. 2017. “Horizontal Gene Transfer Is Not a Hallmark of the Human Genome.” Genome Biology 18 (1): 1–5. https://doi.org/10.1186/s13059-017-1214-2.

Schlötterer, Christian. 2015. “Genes from Scratch--the Evolutionary Fate of de Novo Genes.” Trends in Genetics : TIG 31 (4): 215–19. https://doi.org/10.1016/j.tig.2015.02.007.

Tautz, Diethard, and Tomislav Domazet-Lošo. 2011. “The Evolutionary Origin of Orphan Genes.” Nature Reviews Genetics 12 (10): 692–702. https://doi.org/10.1038/nrg3053.

Vieira-Silva, Sara, and Eduardo P.C. Rocha. 2008. “An Assessment of the Impacts of Molecular Oxygen on the Evolution of Proteomes.” Molecular Biology and Evolution 25 (9): 1931–42. https://doi.org/10.1093/molbev/msn142.

Weibel, Catherine, Jennifer James, Sara M Willis, Paul G Nelson, and Joanna Masel. 2020. “The Protein Domains of Vertebrate Species in Which Selection Is More Effective Have Greater Intrinsic Structural Disorder.” BioRxiv.

Weisman, Caroline M., Andrew W. Murray, and Sean R. Eddy. 2020. “Many, but Not All, Lineage-Specific Genes Can Be Explained by Homology Detection Failure.” PLoS Biology 18 (11): 1– 24. https://doi.org/10.1371/journal.pbio.3000862.

Weiss, Madeline C., Filipa L. Sousa, Natalia Mrnjavac, Sinje Neukirchen, Mayo Roettger, Shijulal Nelson-Sathi, and William F. Martin. 2016. “The Physiology and Habitat of the Last Universal Common Ancestor.” Nature Microbiology 1 (9): 1–8. https://doi.org/10.1038/nmicrobiol.2016.116.

Wherry, R. J. 1931. “A New Formula for Predicting the Shrinkage of the Coefficient of Multiple Correlation.” The Annals of Mathematical Statistics 2 (4): 440–57.

Wilson, Benjamin A., Scott G. Foy, Rafik Neme, and Joanna Masel. 2017. “Young Genes Are Highly Disordered as Predicted by the Preadaptation Hypothesis of de Novo Gene Birth.” Nature Ecology and Evolution 1 (6): 0146. https://doi.org/10.1038/s41559-017-0146.

