## Supplementary figures 1-3 for "Differential retention of Pfam domains creates long-term evolutionary trends"

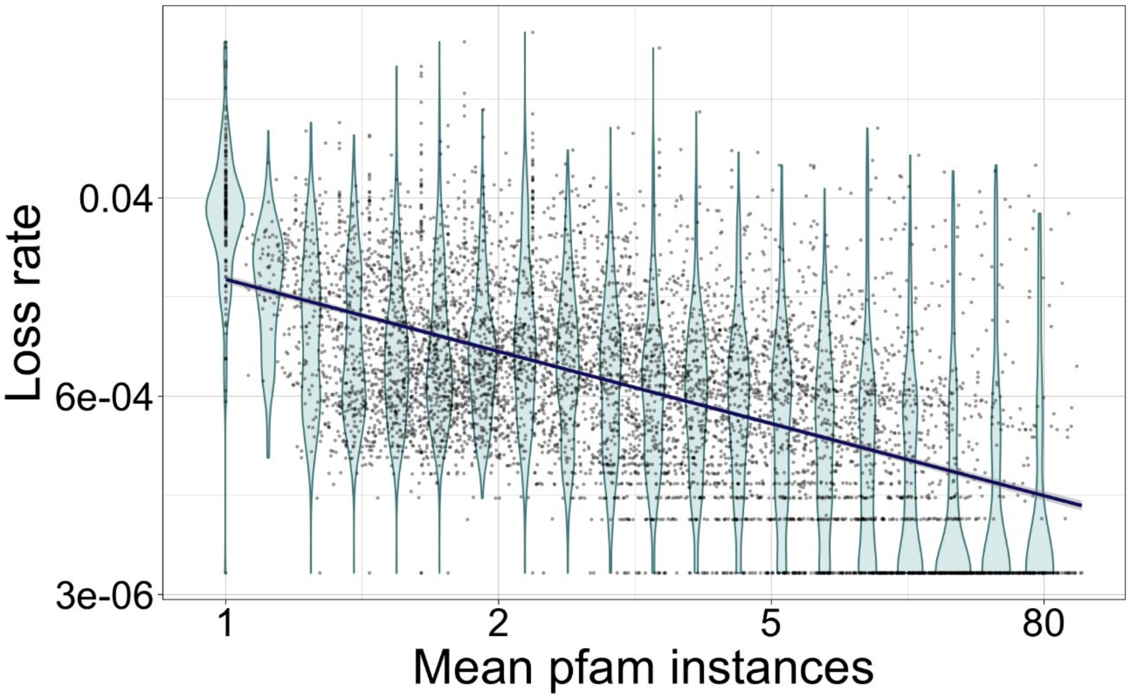


Supplementary figure 1) Pfams with more instances per genome are lost less often. Each point represents a Pfam. Violin plots are a visual guide only, based on dividing the data into 20 groups of equal range of number of Pfam instances. The dark blue line shows the linear regression of loss rate on mean instances, with the 95% confidence intervals shown in grey. Adjusted $R^2 = 0.30$, $p < 1e\text{-}16$.

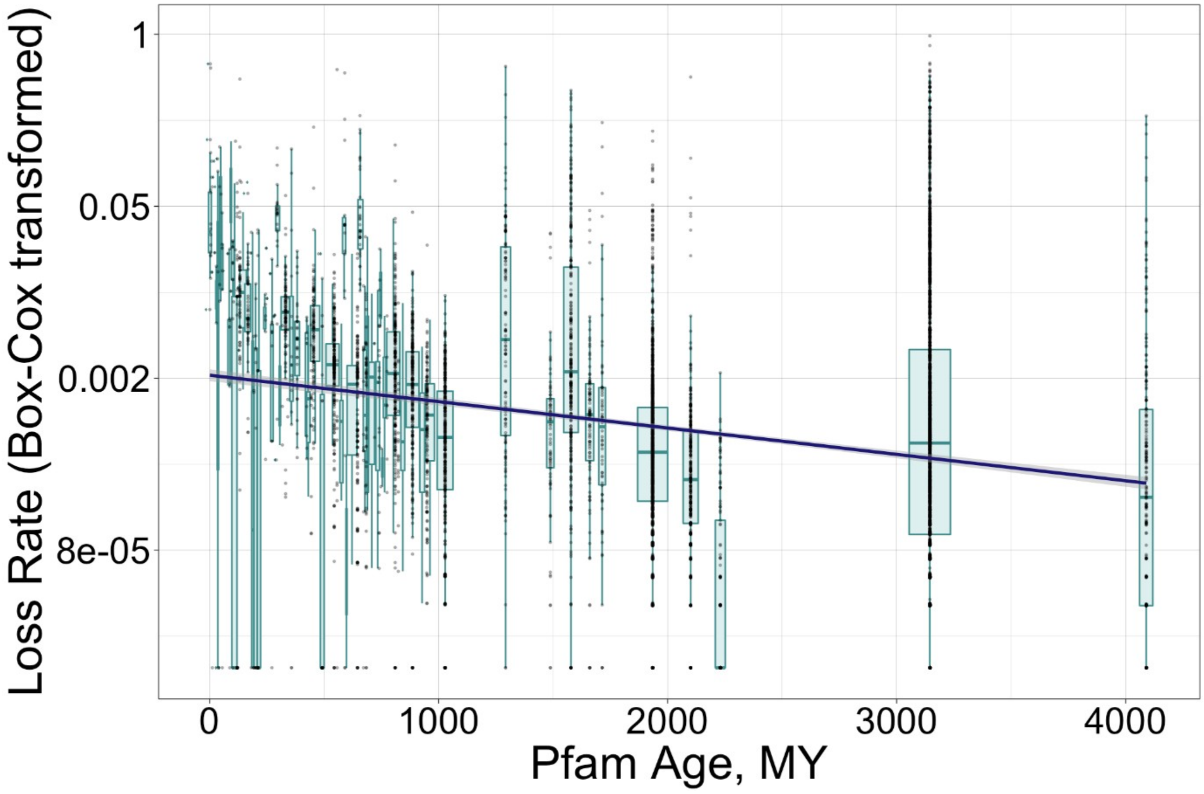


Supplementary figure 2) Older Pfams exhibit lower loss rates. Each box represents all Pfams of a particular age category, with the widths of the boxes proportional to the square root of the number of Pfams in the group. Boxplots show the median, upper, and lower quartiles of the data, while the whiskers represent the largest and smallest data value above and below the interquartile range multiplied by 1.5. Grey points show the values for individual Pfams. The dark blue line is the linear regression slope of the effect of age on loss rate, with the 95% confidence interval shown in grey.

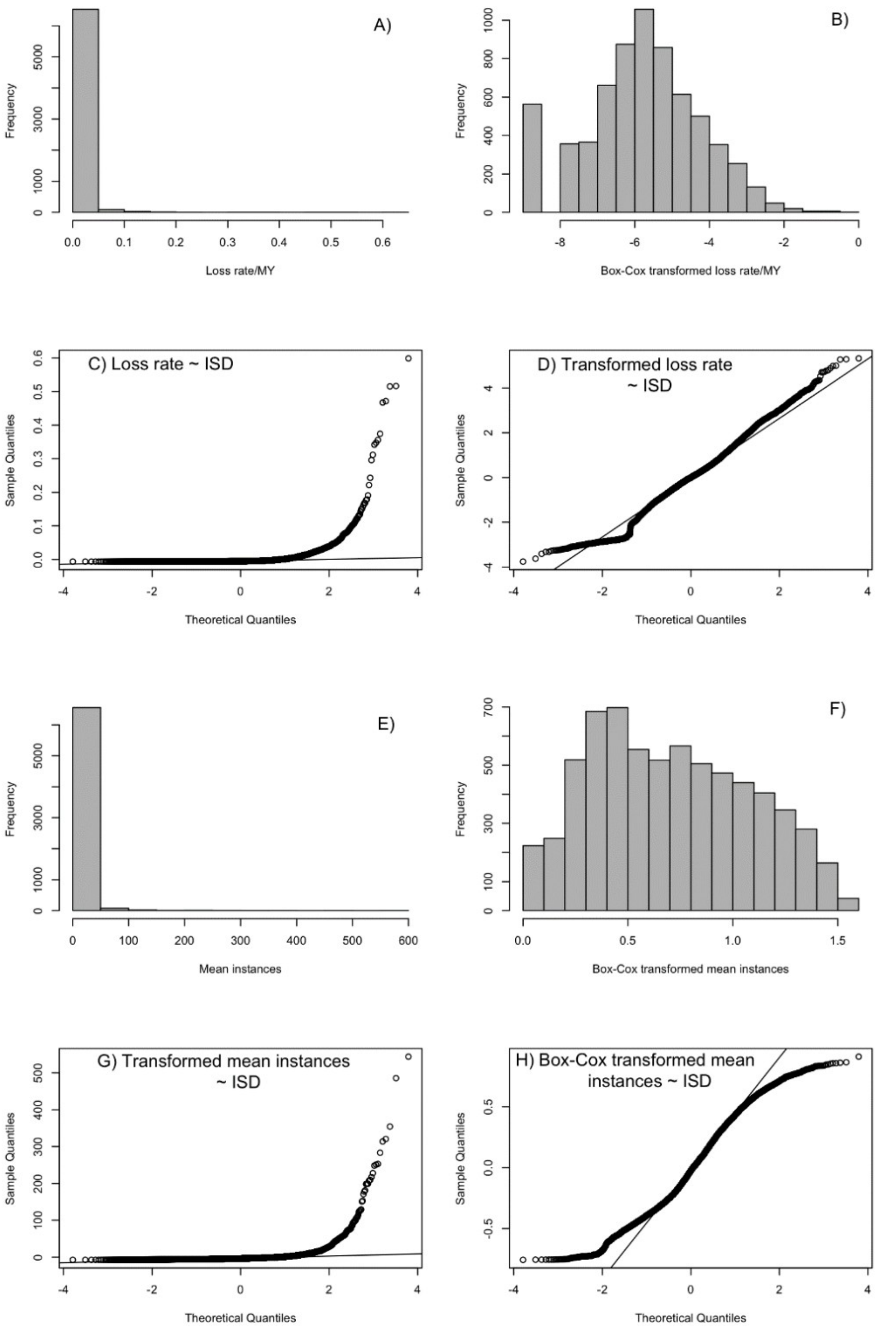


Supplementary figure 3) Transformation of loss rate and mean number of instances is required prior to linear regression. A) and B) show histograms for untransformed and two-parameter Box-Cox transformed ($\lambda = 0.0054$, $\lambda 2 = 6.1e\text{-}06$) inferred loss rates per MY of time. C and D) are Q-Q plots for linear regression models of loss rate and ISD, illustrating how transforming loss rate results in more normally distributed residuals, and is thus more appropriate for our analyses. E) to G) show the corresponding plots for mean Pfam instances (Box-Cox transform $\lambda = 0.058$).
